# Variable rRNA 2’-*O*-methylation fine-tunes ribosome function in *Saccharomyces cerevisiae*

**DOI:** 10.1101/2024.08.07.607065

**Authors:** Sandra Gillot, Virginie Marchand, Yuri Motorin, Agnès Baudin-Baillieu, Olivier Namy

## Abstract

Cellular processes are governed by the regulation of gene expression, often at the level of translation control. The mechanisms of control have been shown to operate at various levels, but there is growing evidence to suggest that rRNA modification patterns play a key role in driving translational modulation of the ribosome. We investigated the intricate relationship between modification status and the decoding activity of the ribosome. We found that the level of 2’-*O*-methylation at specific nucleotides in the rRNA affects the properties of the ribosome, with consequences for both *Saccharomyces. cerevisiae* cell growth and antibiotic sensitivity. More precisely, we demonstrate that methylations within the peptide exit tunnel play an important role in nascent peptide folding. We also demonstrate the modulation of IRES-driven translation by variable methylation at the intersubunit surface of the 60S ribosomal subunit. These findings deepen our understanding of the mechanisms by which 2’-*O*-methylation confers functional specificity on the ribosome.

## Introduction

Translation must be tightly controlled to adapt gene expression to cell requirements. This regulation can be achieved by many factors in the complex translation program, including mRNA properties (1) and abundance (2), translational factors that bind the ribosome (3) and the ribosome itself. Indeed, ribosomes vary due to differences in the sequence of the rDNA alleles expressed (4), ribosomal protein content, posttranslational modifications to these proteins (for review see (5)), and rRNA modifications. Indeed, rRNA is the RNA species displaying the largest numbers of modifications, the most frequent of which being ribose methylation (Nm) and pseudouridine (λϑΙ). The presence of these modifications has long been known, but a major advance was recently made with the discovery of the mechanism by which C/D box guide snoRNAs work with the fibrillarin/Nop1 methyltransferase to control 2’-*O*-methylation at specific positions in rRNA (6, 7). This discovery was followed by the compilation of snoRNA sequences (8, 9), and descriptions of their mode of action and their role in ribosome structure and function (for review see (10)). Indeed, the methylation of 2’-*OH* sites renders nucleotides more hydrophobic, protects against nucleolytic attack, stabilizes helices and increases steric hindrance (11). Nm modifications may interfere with RNA interactions with other RNA molecules or proteins that are dependent on 2′-*OH* groups, thereby enhancing or disrupting inter- or intramolecular interactions (12). The observation that these modifications are conserved in all three kingdoms and positioned within key functional and structural regions of the ribosome led to the consensus view that they must be crucial for the translation process (for a review, see (13)). Many studies have since described the role of particular modifications, individually or in combination, in the decoding capacity of the ribosome (14). However, until recently, it remained challenging to determine the precise level and pattern of rRNA modifications across the entire ribosome population within cells, tissues or organisms. Recent advances in the precise mapping and quantification of these modifications have constituted a major breakthrough. The techniques used are based on deep sequencing (next-generation sequencing or NGS) and provide data with a single-nucleotide resolution. For 2’-*O*-Me detection, RiboMethSeq is the most widely used method (15, 16). RiboMethSeq can be used to map novel potential Nm methylation sites, but it also provides a quantification score reflecting the fraction of such positions modified in the population. Based on these discoveries, it is possible to classify methylation sites into two categories: 1) fully methylated sites that are invariant across cells; and 2) partially methylated sites, implying a certain heterogeneity of the composition of the ribosome pool either within or between cells (17). This observation paved the way for exploration of the methylation status of ribosomes in various organisms as a function of developmental stage (18–20) or disease (21, 22). The characterization of hypo-2’- *O*-methylated ribosomes resulting from deletion of the yeast Hit1 and Bcd1 proteins involved in snoRNP biogenesis and snoRNA stability (23, 24) and studies of fibrillarin knockdown in human cells (25) have revealed considerable variability in the 2’-*O*-methylation patterns of rRNA, with consequences for translation. Indeed, following the use of siRNAs decreasing fibrillarin protein levels five-fold in human HeLa cell lines, almost all the 106 2’-*O*-methylated positions identified displayed decreases in methylation that were highly variable (between 0.2 and 57%) and highly position-specific (25).

We used *S. cerevisiae* as a model and focused on the subset of conserved 2’-*O*-Me sites displaying the largest decreases in fibrillarin levels in HeLa cells. We reasoned that the cell might make use of the dynamics of rRNA methylation to adapt its translational program to changing environmental conditions. We therefore systematically tested the translational accuracy of ribosomes by altering the pattern of Nm methylations close to functional regions. We found that the plasticity of rRNA 2’-*O*-methylation could control the differential translation of mRNA by conferring decoding specificity on the ribosome.

## Results

### Engineering of yeast *S. cerevisiae* strains with alterations to rRNA modification

For the construction of yeast ribosomes with an altered methylation status, we began by considering the 32 rRNA 2’-*O*-methylation sites identified in the *H. sapiens* ribosome as strongly affected (decrease of more than 17%) by fibrillarin depletion (25). The yeast ribosome contains 55 known 2’-*O*-methylation sites, many of which are located in positions conserved between yeast and humans. We therefore performed a comparative analysis of Nm residues in yeast and human ribosomes (8, 9). Only nine of the 32 highly variable Nm positions in humans were conserved in the yeast *S. cerevisiae* (see Table 1). We visualized their locations in the 3D structure of the yeast ribosome and clustered them into six main regions, highlighted in different colours in Figure 1, which, as expected, mostly mapped to functional regions of the ribosome. Based on the degree of conservation and/or the number of positions within a particular domain, we decided to focus on three major functional domains: i) the E-site; ii) the peptide exit tunnel (PET) and iii) the intersubunit surface of 60S (Table 1).

**Figure 1.**
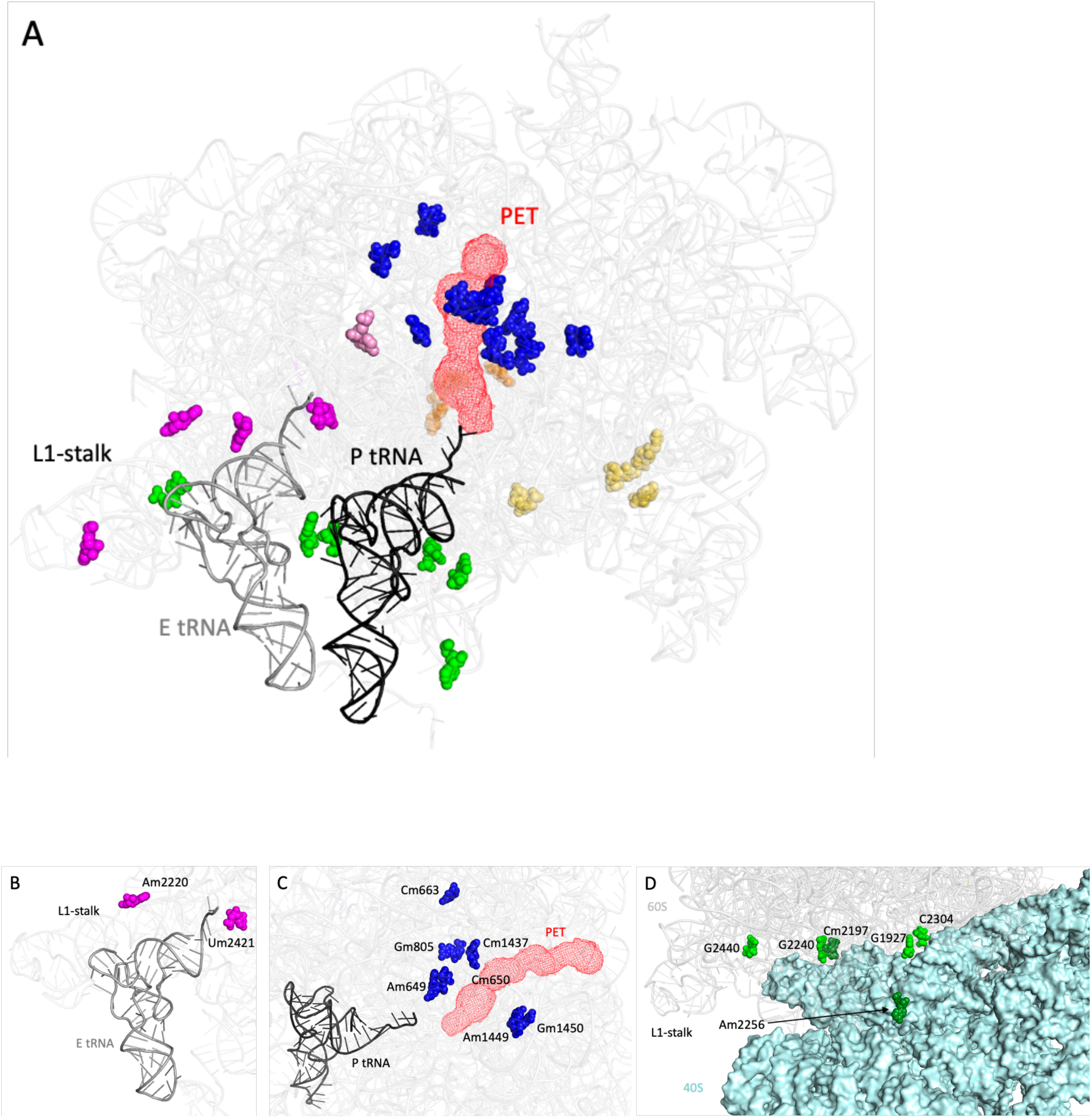
Position and color-coded classification of the rRNA 2’-*O*-methylated nucleotides studied on the 3D structure of the *S. cerevisiae* 60S subunit. (*A*) The position of each human hypo-2-*O*-methylated site within the 5.8S and 25S rRNA subunits (Table 1) is reported on the yeast 80S structure (PDB 3J78) and color-coded according to its location in functional domains (E-site in magenta, peptide exit tunnel in blue and intersubunit surface in green) or uncharacterized domains (spread beneath the surface in yellow, clustered within an internal pocket in orange and within an internal protein-free pocket in pink). (*B*-*D*) View of three functional domains showing the positions in yeast that are affected by the loss of snoRNAs within the E-site (*B*) and the peptide exit tunnel (*C*) or by the expression of an engineered snoRNA (light green) at the intersubunit surface (*D*).

**Table 1.**
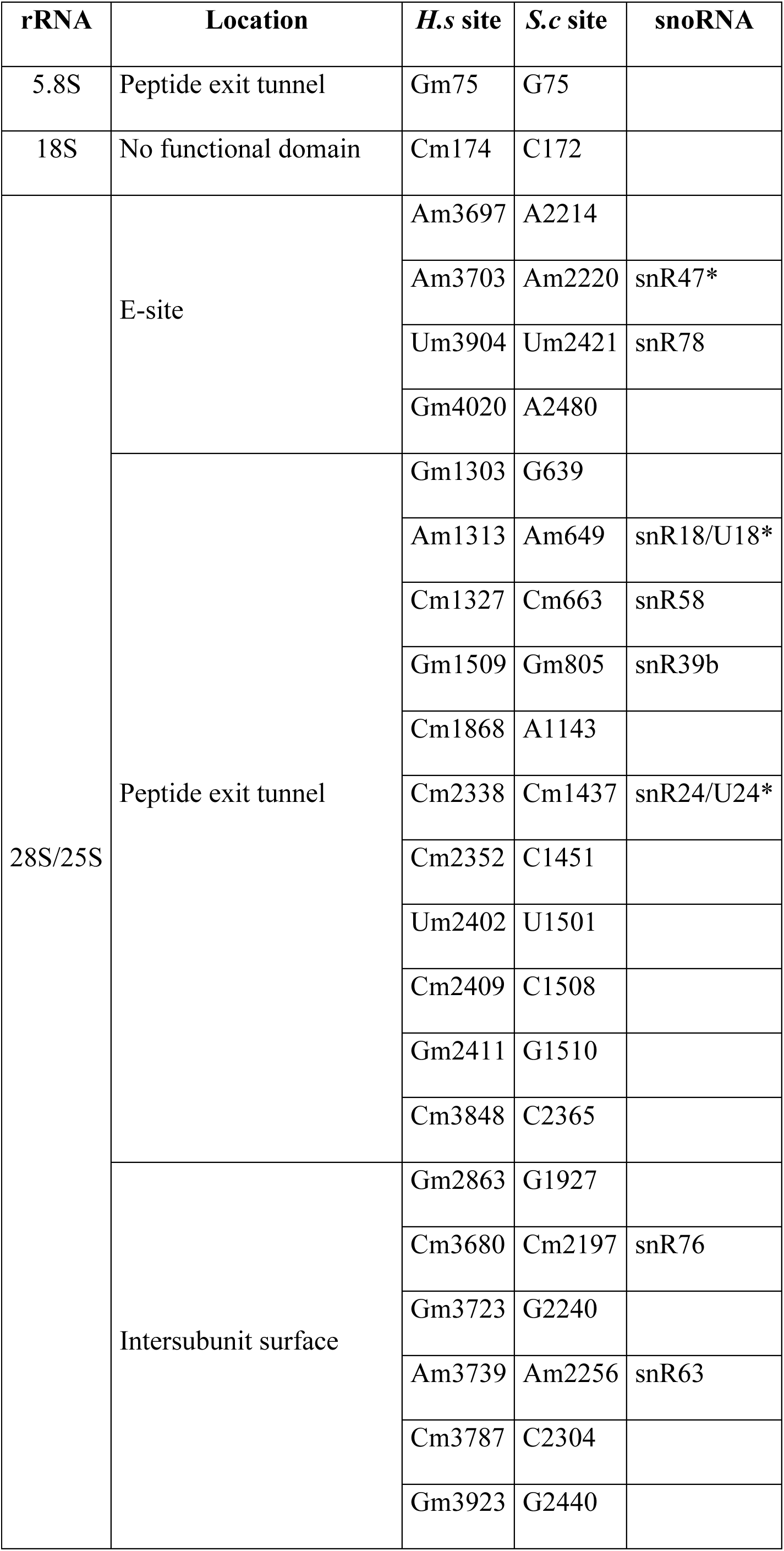
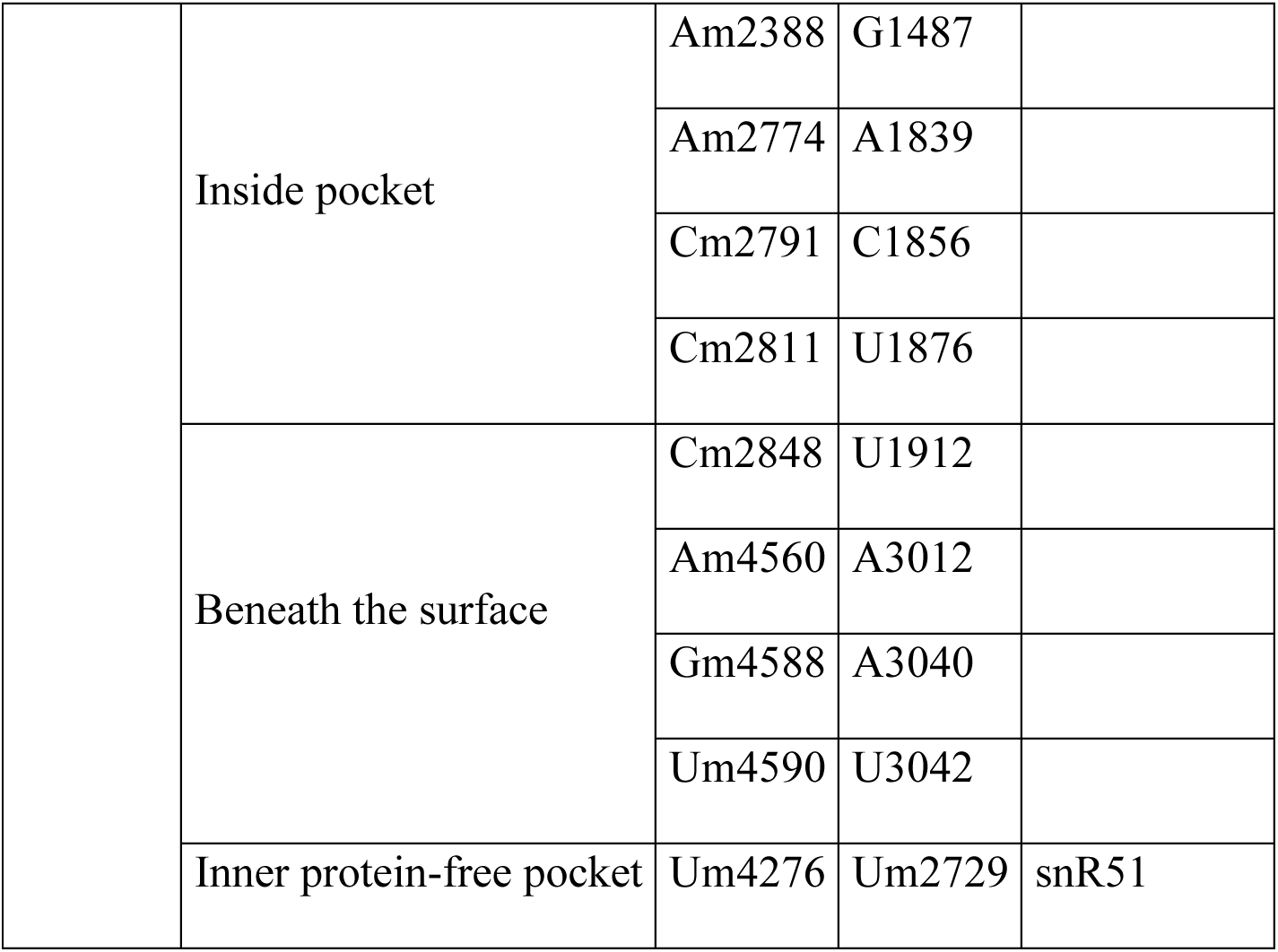
Correspondence between human and yeast variable 2’-*O*-Me sites. Human rRNA nucleotides hypo-2’-*O*-methylated under conditions of fibrillarin depletion (threshold above 17%) are listed and classified according to their location within the ribosome. The corresponding *S. cerevisiae* rRNA nucleotides are given. The snoRNAs are indicated for the nine modified positions conserved in yeast. * indicates that this snoRNA also directs methylation at additional sites (see text).

| **rRNA** | **Location** | ***H.s* site** | ***S.c* site** | **snoRNA** |
| --- | --- | --- | --- | --- |
| 5.8S | Peptide exit tunnel | Gm75 | G75 |  |
| 18S | No functional domain | Cm174 | C172 |  |
| 28S/25S | E-site | Am3697 | A2214 |  |
|  |  | Am3703 | Am2220 | snR47* |
|  |  | Um3904 | Um2421 | snR78 |
|  |  | Gm4020 | A2480 |  |
|  | Peptide exit tunnel | Gm1303 | G639 |  |
|  |  | Am1313 | Am649 | snR18/U18* |
|  |  | Cm1327 | Cm663 | snR58 |
|  |  | Gm1509 | Gm805 | snR39b |
|  |  | Cm1868 | A1143 |  |
|  |  | Cm2338 | Cm1437 | snR24/U24* |
|  |  | Cm2352 | C1451 |  |
|  |  | Um2402 | U1501 |  |
|  |  | Cm2409 | C1508 |  |
|  |  | Gm2411 | G1510 |  |
|  |  | Cm3848 | C2365 |  |
|  | Intersubunit surface | Gm2863 | G1927 |  |
|  |  | Cm3680 | Cm2197 | snR76 |
|  |  | Gm3723 | G2240 |  |
|  |  | Am3739 | Am2256 | snR63 |
|  |  | Cm3787 | C2304 |  |
|  |  | Gm3923 | G2440 |  |
|  | Inside pocket | Am2388 | G1487 |  |
|  |  | Am2774 | A1839 |  |
|  |  | Cm2791 | C1856 |  |
|  |  | Cm2811 | U1876 |  |
|  | Beneath the surface | Cm2848 | U1912 |  |
|  |  | Am4560 | A3012 |  |
|  |  | Gm4588 | A3040 |  |
|  |  | Um4590 | U3042 |  |
|  | Inner protein-free pocket | Um4276 | Um2729 | snR51 |

In human ribosomes, four Nm residues close to the E-site (28S-Am3697/28S-Am3703/28S-Um3904 and 28S-Gm4020) are affected by fibrillarin depletion. Two of these Nm residues are conserved in *S. cerevisiae* (25S-Am2220 and 25S-Um2421, shown in magenta in Fig. 1*A* and *B*). They are targeted by C/D-box snoRNAs encoded by the monocistronic genes snR47 and snR78, respectively. This simple genomic organization facilitated the construction of *S. cerevisiae* strains with deletions of single or multiple snoRNA genes, termed Δ2^E^ (Table 2).

**Table 2.**
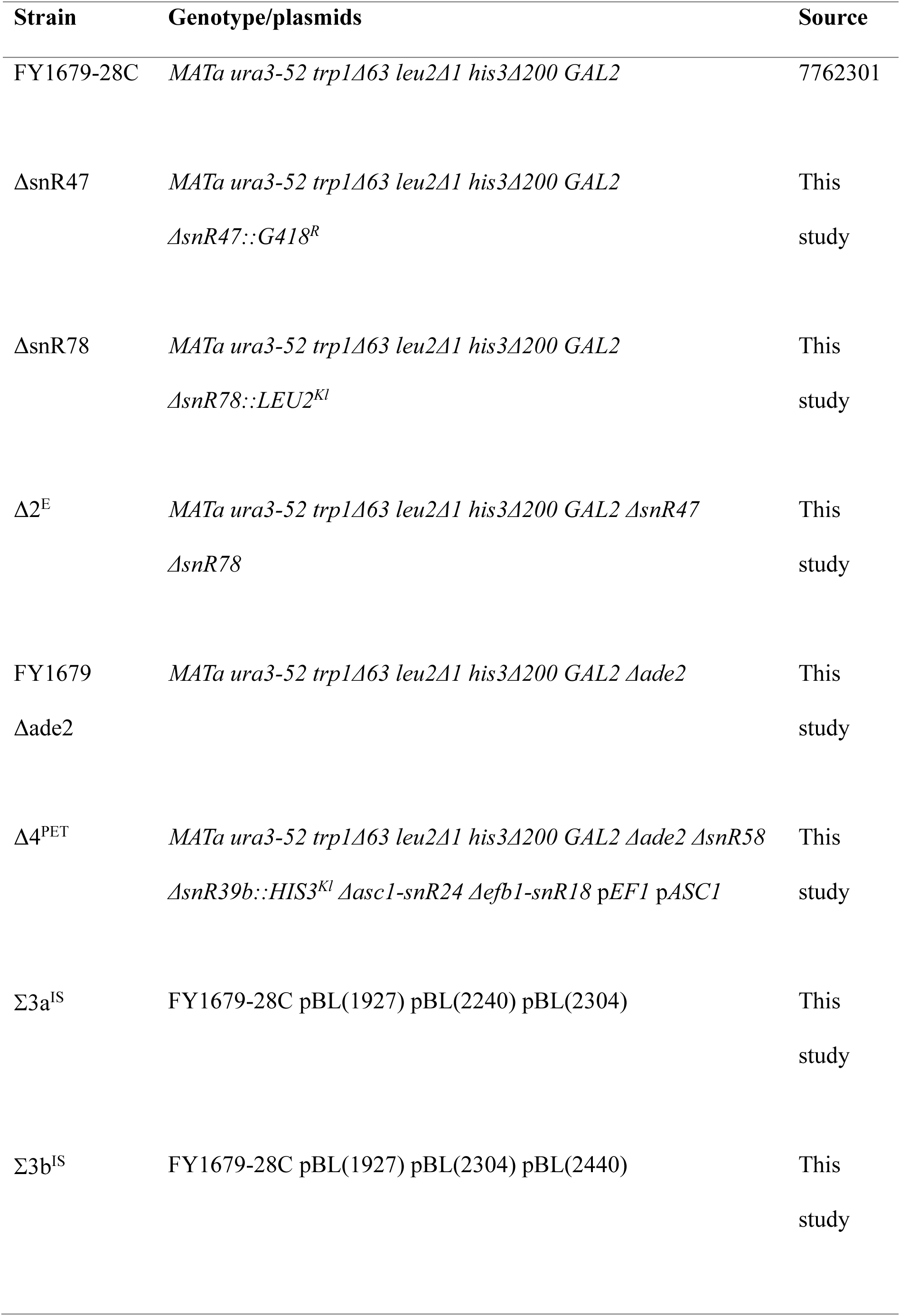
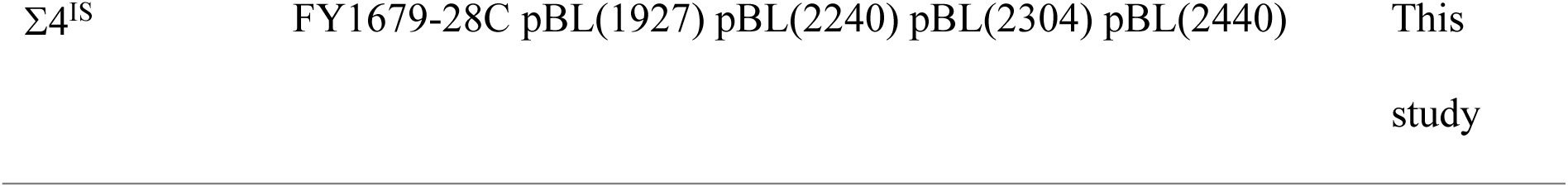
List of yeast strains investigated in this study.

As many as 12 Nm modifications are clustered around the PET of the human ribosome, 11 on the 28/25S rRNA and one on the 5.8S rRNA (shown in blue in Fig. 1*A* and *C*). Four of these sites are conserved between human and yeast ribosomes: 25S-Am649 targeted by snR18/U18, 25S-Gm805 targeted by snR39b, 25S-Cm663 targeted by snR58 and 25S-Cm1437 targeted by snR24/U24. The study of this set of modifications is complicated by the location of snR18/U18 and snR24/U24 within the intronic regions of *EFB1* (26) and *ASC1* (27), respectively. These two genes encode proteins involved in translation. Moreover, those two snoRNAs also guide additional methylations: snR18/U18 targets the adjacent position 25S-Cm650 specifically in yeast and snR24/U24 also targets the 25S-Am1449 and 25S-Gm1450 positions located within the PET. We addressed the question of the global effects of a loss of rRNA methylation within the PET by constructing a yeast strain in which all four snoRNAs were deleted (denoted Δ4^PET^) complemented by two plasmids expressing the open reading frames of *EFB1* and *ASC1,* which therefore lacked the seven 2’-*O*-methylation modifications mentioned above.

Finally, six modified positions on the intersubunit surface of 60S displayed a substantial decrease in their methylation status following fibrillarin depletion in human cells (shown in green in Fig. 1*A* and *D*). Only two of these positions are conserved in yeast — 25S-Cm2197 (guided by snRNA76) and 25S-Am2256 (guided by snR63) — and deletion of the two corresponding snoRNAs to abolish these Nm methylations would therefore provide only partial information about the importance of these six modifications in humans. We instead introduced the four missing methylations (at 25S-G2240, 25S-G1927, 25S-G2440 and 25S-C2304) into yeast rRNA via an *in vivo* site-directed methylation strategy (27, 28). This approach is based on the hijacking of naturally encoded snoRNAs for the specific methylation of selected nucleotides in the rRNA through the replacement of the natural snoRNA guide element by a customized guide sequence. The nucleotide to be modified is targeted through specific base pairing of the snoRNA guide sequence to the rRNA substrate. We generated snR38-derived sequences specifically targeting positions G1927, G2240, C2304 and G2440 in yeast 25S rRNA, which we introduced into wild type (WT) *S. cerevisiae* cells.

### Loss of methylation within the E site or the PET has little or no effect on growth

All the yeast *S. cerevisiae* strains expressing ribosomes with deficient rRNA 2’-*O*-methylation in the aforementioned regions generated were viable on standard rich media (see Mat&Meth). We therefore explored the growth and functional defects of these strains in more detail.

Strains bearing deletions of snR47 and snR78 targeting the E-site Nm residues either individually (data not shown) or in combination (ι12^E^) were affected only slightly if at all in terms of growth, ribosomal biogenesis and/or global translation rates, as indicated by their normal polysome levels of 40S, 60S and 80S (Fig. S1*D*).

The ι14^PET^ mutant strain lacking the four snoRNAs targeting the PET grew slightly less well than the wild type, with a 38% longer doubling time (4.7 h versus 3.4 h, Fig. 2A). We therefore also assessed the effects of this quadruple deletion on ribosome biogenesis and global translation efficiency by polysome profiling, in which mRNAs are separated on a sucrose gradient according to the number of bound ribosomes, providing an accurate reflection of the translation status of the cell. The absence of the four snoRNAs led to a slight deficit of 60S and the presence of half-mers reflecting the presence of the 40S subunit and its inability to combine with the absent 60S to form an 80S functional ribosome (Fig. S1*A* and *B*). Such a phenotype was previously described for the loss of snR24 (29). We, thus, decided to complement the ι14^PET^ mutant strain with snR24 expressed from the intron of the yeast actin gene (6). The expression of snR24 alone completely restored 60S subunit levels and abolished the defects in polysome profiling, demonstrating that these rRNA modifications in the PET play only a minor role in ribosome biogenesis (Fig. S1*C*). We observed no marked decrease in the number of translating ribosomes, but the lengthening of the generation time in the mutant strain suggested that Nm methylation in the PET was important for translation for the efficient maintenance of cellular homeostasis.

**Figure 2.**
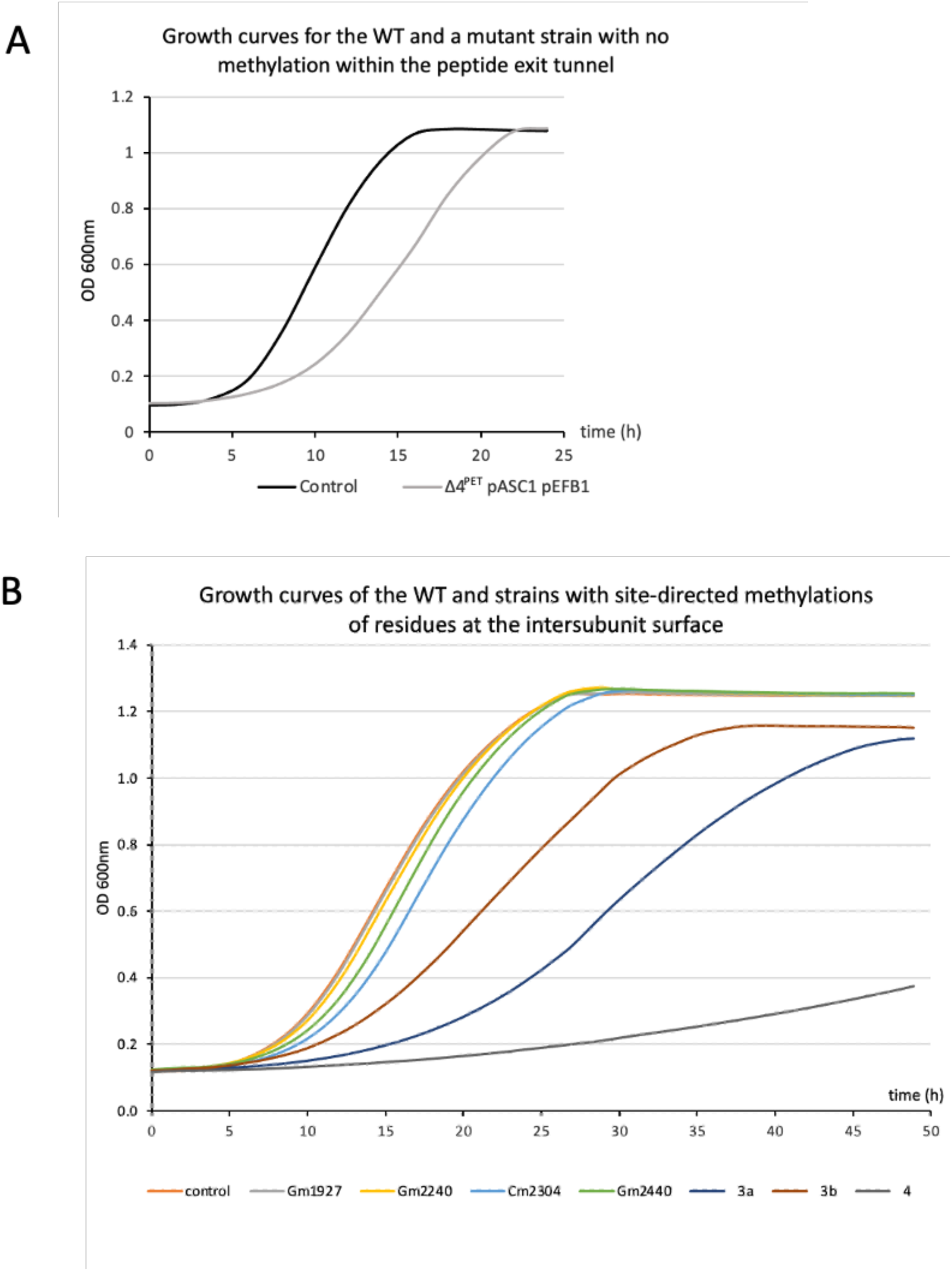
Strains with a modified methylation status in the peptide exit tunnel or the intersubunit surface have a slow-growth phenotype. (*A*) Yeast cells from control and 1′4^PET^ strains were cultured in 250 ml minimal SD glucose medium to select for plasmids expressing the *ASC1* and *EFB1* open reading frames. (*B*) Control yeast cells were transformed with the pBL152 plasmid encoding the natural snoRNA snR38 targeting G2811 as a negative control or a combination of pBL vectors introducing additional modifications at the intersubunit surface and cultured at 30°C in 250 ml minimal SD galactose medium to induce site-directed methylation. Strains were cultured for 24 h to 48 h, at 30°C in a TECAN apparatus, with hourly measurements of OD_600_.

### Additional “human-like” Nm modifications introduced into yeast rRNA substantially compromise growth and translation

We first checked that the artificial snoRNAs expressed in the WT strain of *S. cerevisiae* guided rRNA modifications at the expected locations. To that purpose, we performed RiboMethSeq analysis (15), which measures the protection of the RNA phosphodiester bond against cleavage under alkaline conditions. The methyl group at the 2’-*OH* of the ribose efficiently protects RNA from cleavage, making it possible to detect any new “alkali-resistant” locations. Total RNA extracted from yeast strains grown on galactose to induce custom snoRNA extraction was subjected to RiboMethSeq analysis. We checked that *de novo* methylation at the four expected positions was efficient and without off-target effects, by inspecting the MethScore obtained for the presumed positions of modifications and for all rRNA positions. We found that snoRNA-directed methylation was efficient for all additional positions in the *S. cerevisiae* 25S rRNA, with no off-target methylation at other sites (Fig. S2).

An analysis of this engineered yeast strain demonstrated that the introduction of additional “human-like” Nm modifications to the 60S intersubunit surface had a much stronger effect than the other mutations considered. A single additional methylation had almost no impact on growth (Fig. 2*B*), but the addition of three extra rRNA modifications (Σ3a^IS^ or Σ3b^IS^) significantly decreased growth, and growth was almost entirely abolished by the combination of all four additional Nm (Σ4^IS^) (Fig. 2*B*). We therefore decided to investigate global translation levels in this artificial *S. cerevisiae* strain. We first performed polysome profiling to calculate the polysome-to-monosome ratio (P/M) (Fig. 3*A*). The P/M ratio of the control strain was 1.07, which was significantly higher than the ratio for the Σ4^IS^ strain, which was 0.68. This decrease in P/M ratio indicates a decrease in the overall rate of translation. For confirmation of this observation, we used the Click-iT™ HPG Alexa Fluor™ 488 Protein Synthesis Assay to quantify nascent protein synthesis. This method uses the noncanonical amino acid L-homopropargylglycine (HPG), which is incorporated into proteins in place of methionine. Strains were grown on galactose to induce additional “human-like” Nm methylation and the incorporation of HPG was followed for 60 min (Fig. 3*B*). A clear decrease in HPG incorporation was observed at all time points for the hypermethylated strain. These data confirm the significantly lower rate of translation in the Σ4^IS^ strain than in wild-type yeast.

**Figure 3.**
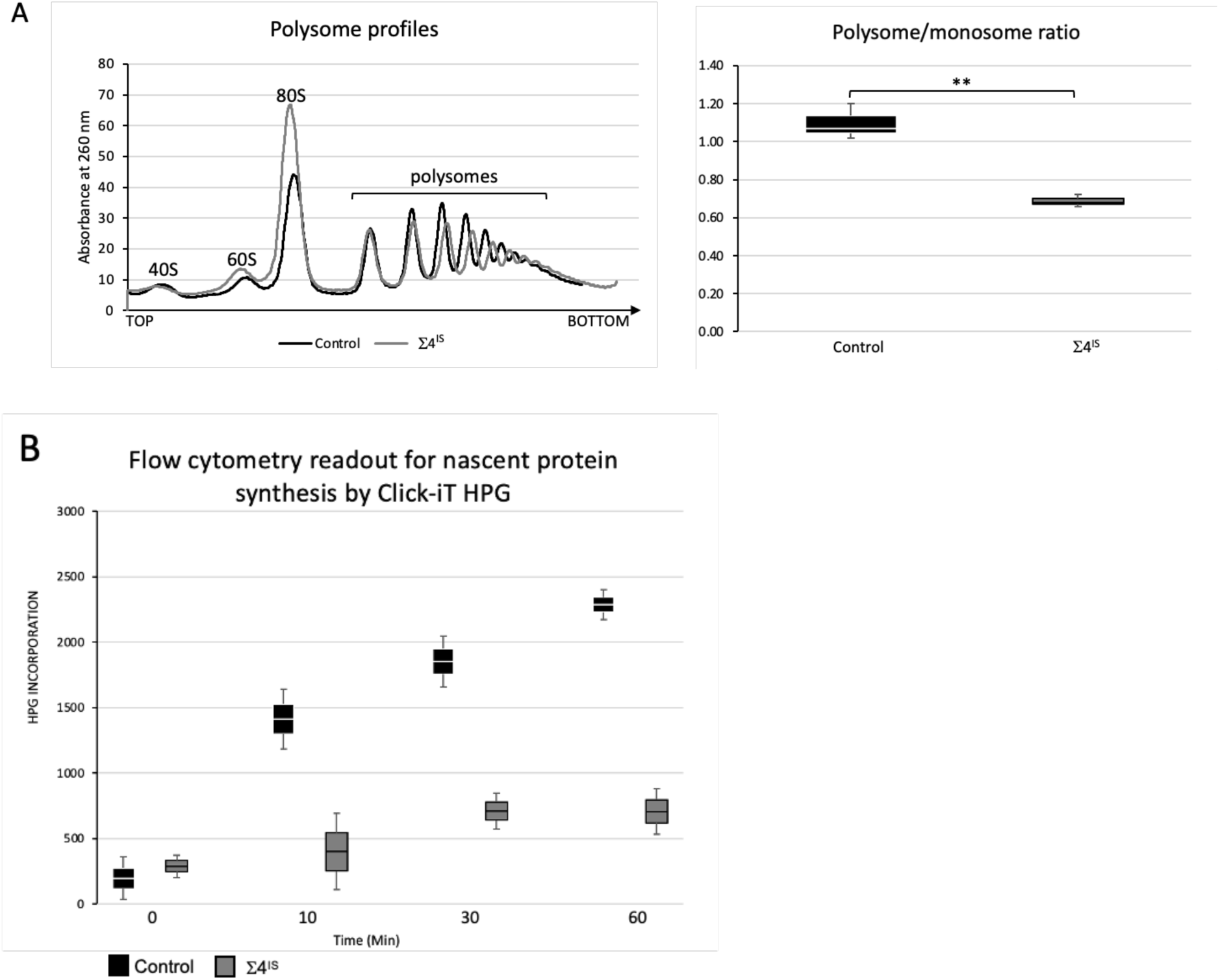
Global translation is affected by additional methylation at the intersubunit surface. (*A*) Polysome profiles and boxplot presenting polysome-to-monosome ratios for control and Σ4^IS^ strains grown in the presence of galactose. The corresponding peaks (80S/monosomes and polysomes) are indicated on the polysome profile. P/M ratios were calculated with ImageJ from the areas beneath the profile curve corresponding to the monosome (80S) and polysome regions (3 experiments, ** *p*=0.01) (*B*) Incorporation of HPG into newly synthesized proteins in control and Σ4^IS^ cells, assessed with the Click-iT kit.

### Missing or additional Nm methylations have a specific effect on the decoding capacities of the ribosome

We used a dual reporter system based on the relative levels of expression of two reporter genes to investigate whether missing Nm sites or the introduction of extra Nm sites in yeast ribosomes affected translational accuracy. In the constructs used, the coding sequences of β-galactosidase and firefly luciferase encode a translational fusion but are separated by recoding sequences that induce either frameshifts or stop codon readthrough (Fig. 4*A*). We used these constructs to assess the accuracy of the translation elongation and termination steps with sequences: 1) known to promote -1 (BLV or IBV) or +1 (EST3) frameshifting, and 2) for termination, a readthrough sequence (TMV) containing an in-frame UAG stop codon, and when required, the UAA and UGA stop codons (see Experimental Procedures).

**Figure 4.**
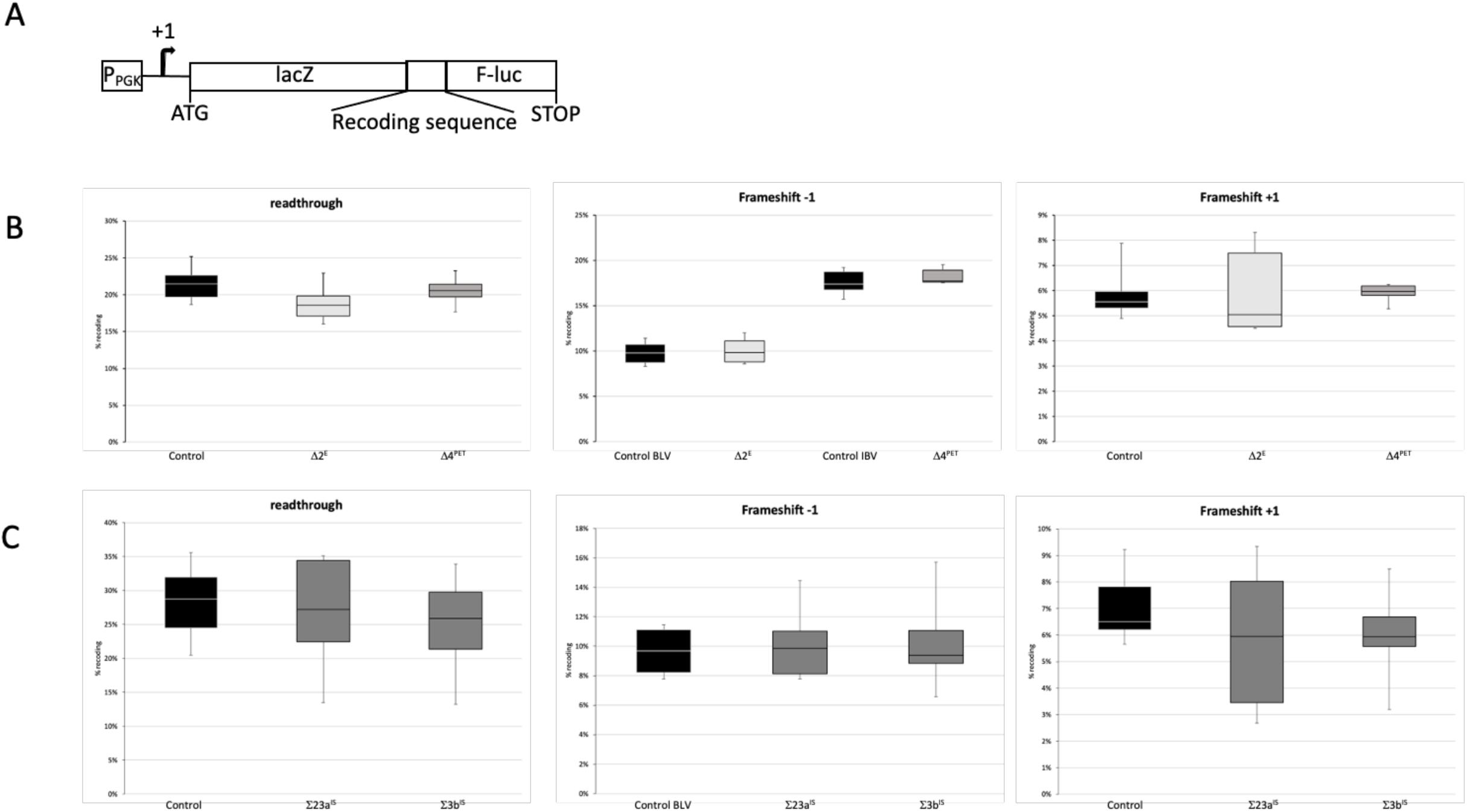
Alterations to methylation status do not impair translation fidelity or stop codon recognition efficiency. (*A*) Structure of the reporter gene. A sequence containing a programmed recoding event, such as stop codon readthrough (UAG) or -1/+1 frameshifting (BLV/IBV (-1) and EST3 (+1) sequences), was inserted between the *lacZ* and firefly luciferase coding sequences. The fusion gene was expressed under the control of the PGK1 gene promoter. (*B*) Recoding efficiency in the wild type (CTRL) and in mutant strains lacking modifications within the E site (1′2^E^) or the PET (1′4^PET^); (*C*) as in *B* for the wild-type (CTRL) and mutant strains, but with additional modifications of the intersubunit surface and culture in the presence of galactose (Σ3a^IS^ and Σ3b^IS^).

The error rates for each mutant strain lacking methylation at either position were similar to those obtained for the wild-type strain, whatever the recoding event measured (Fig. 4*B*). Similarly, when a combination of 3 methylations at the 60S intersubunit surface was added, translation accuracy was not significantly affected (Fig. 4*C*). Thus, these positions susceptible to variation in the context of fibrillarin depletion are probably not involved in reading-frame maintenance or translation termination efficiency.

Another non-canonical event potentially affected by the missing or extra Nm residues is cap and polyA-independent translation using internal ribosome entry sites (IRES). Initiation at IRES sequences is a complex multistep process. Briefly, the IRES binds to the 40S surface. This binding triggers several conformational changes, allowing the recruitment of the 60S subunit to form an initiation-competent 80S ribosome. Moreover, direct recruitment of the 80S ribosome by the IRES is also possible although disfavored (30). In the resulting 80S–IRES ribosomes, IRES occupies the intersubunit space spanning from the A- to E-sites, where it interacts with the phylogenetically conserved 80S core. As the positions studied are located within these regions, we investigated the initiation of IRES-dependent translation in strains with alterations to the Nm status of the E-site and intersubunit surface. The CrPV IRES is a class IV IRES that enables ribosomes to be recruited to the mRNA without the need for eukaryotic initiation factors or initiator tRNA^Met^. Class IV IRES function in many different species and support translation initiation in the yeast *Saccharomyces cerevisiae* (31), making it possible to use a dual reporter system based on the inclusion in the same mRNA of the cap-dependent *Renilla* luciferase and IRES-dependent firefly luciferase sequences (Fig. 5*A*). For E-site Nm nucleotides, a small non-significant increase in CrPV IRES initiation rates was observed in the absence of snR78 (guiding 25S-Am2220), whereas no effect was detected in the i¢lsnR47 strain (lacking 25S-Um2421) (Fig. 5*B*). These results indicate that neither of these two modifications influence the capacity of the yeast ribosome to initiate translation at such heterologous IRES sites. We then investigated the effect of extra Nm methylations at the 60S intersubunit surface. CrPV-mediated translation was substantially improved by both combinations of additional Nm residues, with rates of 147% and 155%, respectively, relative to the control (Fig. 5*B*). No significant differences were observed when strains were grown on glucose, in conditions in which no additional Nm methylation can occur (data not shown). These results suggest that additional Nm modifications in the human ribosome enhance CrPV IRES-mediated translation initiation.

**Figure 5.**
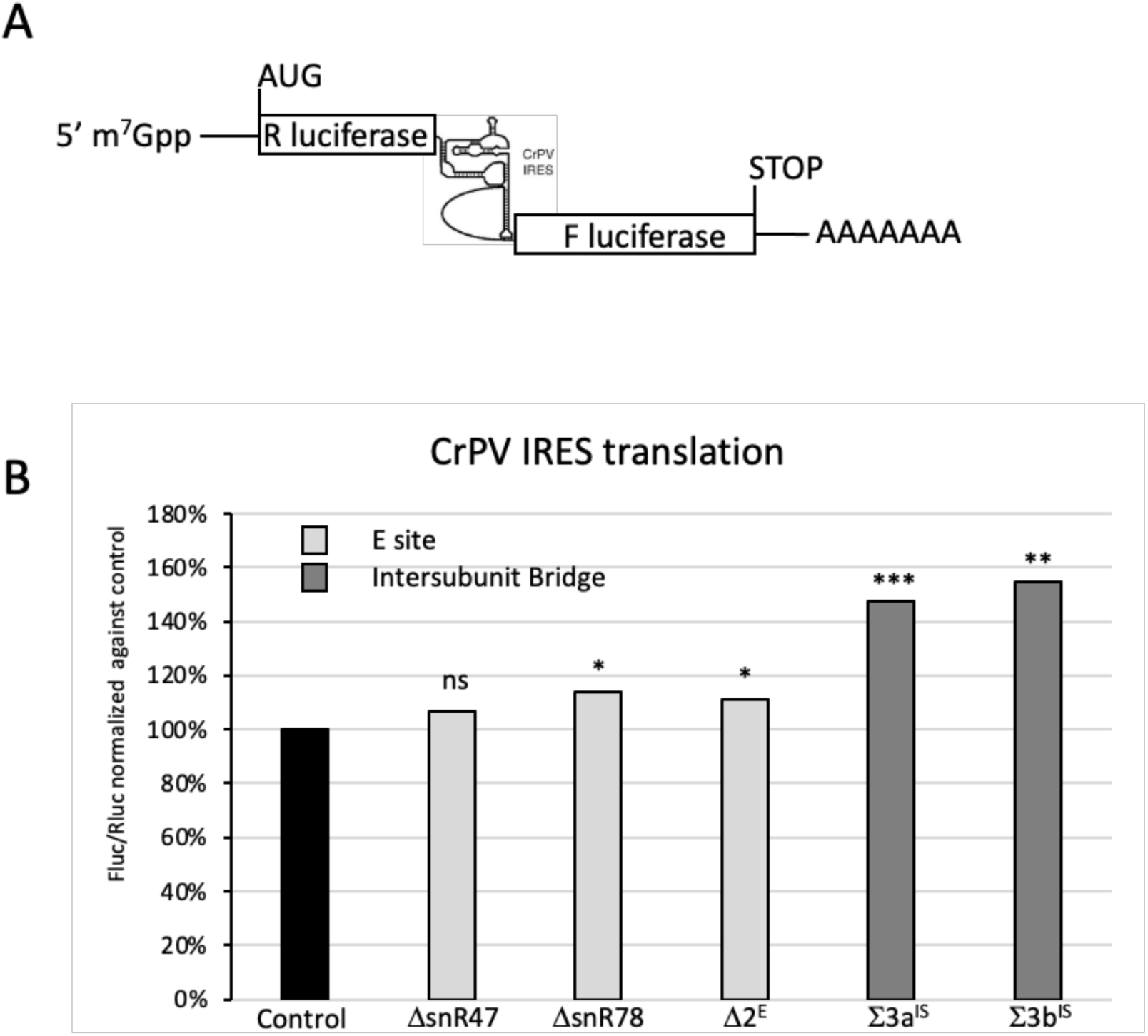
Efficiency of IRES initiation, as determined by calculating the ratio of firefly and *Renilla* luciferase activities. (*A*) Schematic diagram of the reporter system. The fusion gene was composed of the coding sequence of the *Renilla* luciferase, the translation of which is cap-dependent, and the firefly luciferase, the translation of which is IRES-dependent (CrPV). (*B*) The value indicated is the median Fluc/Rluc ratio (*n*=5 to 12) in the hypo- (E site) or hyper- (intersubunit surface) methylated state, normalized against the Fluc/Rluc ratio measured in control cells and expressed as a %. Statistical significance was determined by calculating the *p*–value in a *t*-test (*p*<0.1=*; *p*<0.01=**; *p*<0.001=***; ns=non significant).

### Antibiotic susceptibility is dependent on the location of the modifications

Antibiotic binding to the specific sites in the ribosome can be modulated by rRNA modifications (32–35). We therefore explored the effects of this loss of Nm methylation on resistance/susceptibility to a set of antibiotics binding to various functional centers of the ribosome. In particular, cycloheximide binds to the E-site and blocks the elongation step of translation. Marked susceptibility to cycloheximide was observed in strains lacking 25S-Um2421 (guided by snR78), whereas only a slight effect was observed for the loss of 25S-Am2220 and/or 18S-Am619 (ιβsnR47) (Fig. 6*A*). The Nm of U2421 is, therefore, involved in the resistance of the yeast ribosome to 0.5 μg/ml cycloheximide.

**Figure 6.**
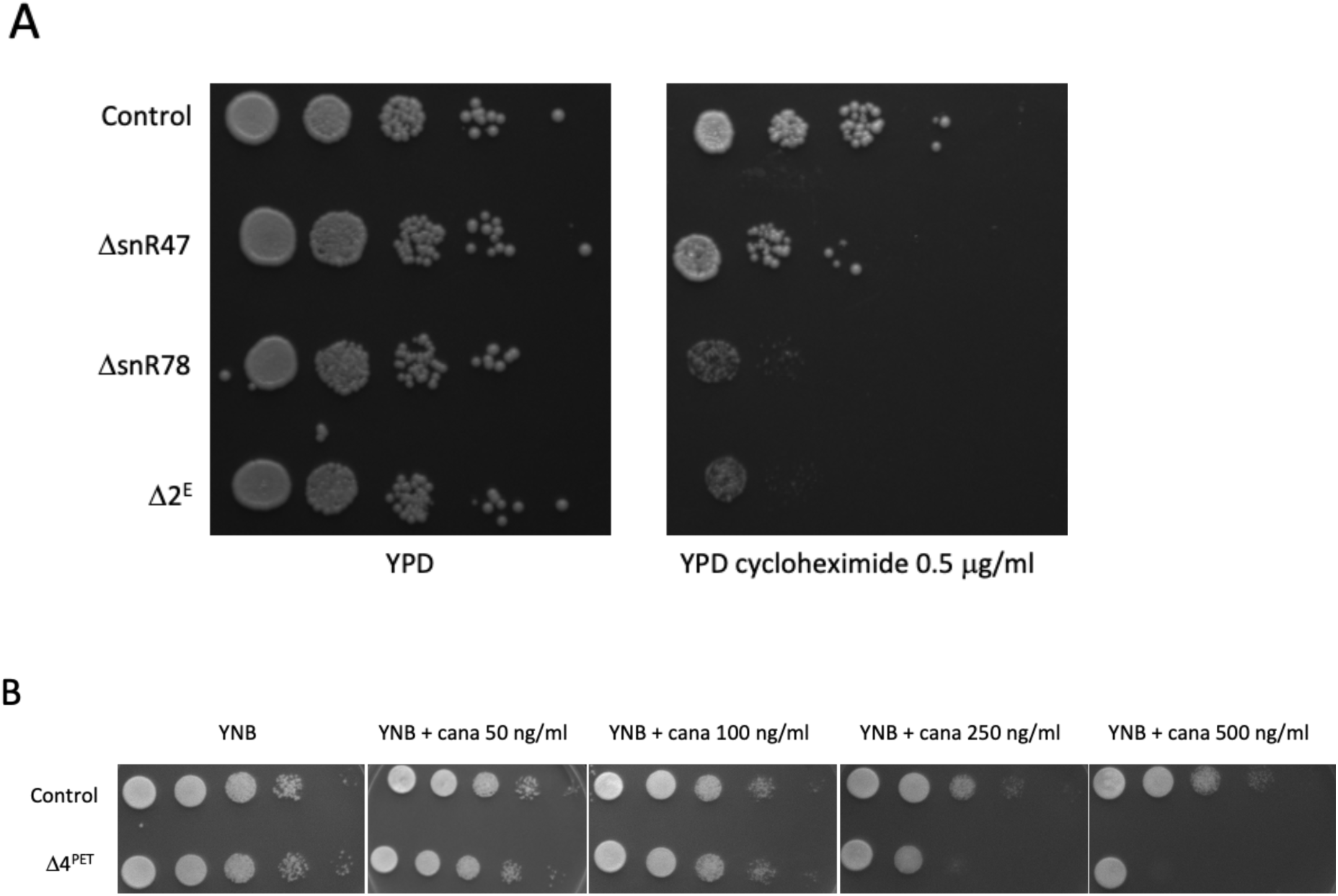
Antibiotic susceptibility tests. (*A*) 1′snR78 mutant cells lacking E-site methylation were more susceptible than the wild type to cycloheximide. 10-fold dilutions of control (FY1679), 1′snR47, 1′snR78 and 1′2^E^ cells were plated on YPD alone or YPD containing 0.5 μg/ml cycloheximide and cultured for three days at 30°C. (*B*) Strain devoid of PET methylations displayed enhanced susceptibility to canavanine. 10-fold dilutions of control (FY1679) and 1′4^PET^ cells were plated on minimal medium (YNB) alone or on YNB supplemented with various concentrations of canavanine (ranging from 50 to 500 ng/ml) and cultured for three days at 30°C.

The PET of the ribosome plays a key role in protein synthesis by serving as a conduit for the emerging polypeptide chain. Beyond its transport function, the PET is involved in quality control, ensuring correct protein folding (36). We investigated the effects of an absence of Nm methylations close to the PET on the function of this structure. For this purpose, we used L-canavanine, a non-proteinogenic amino acid, the incorporation of which into proteins results in aberrant folding (review in (37)). The lack of Nm in the Δ4^PET^ strain greatly decreased the resistance of the strain to this toxic amino-acid analog (Fig. 6*B*). Thus, these 2’-*O*-methylation sites close to the PET contribute to correct protein folding albeit through an as yet unknown mechanism and at a level yet to be determined.

### Modifications within the PET are essential for the correct folding of the nascent peptide

The role of the PET in protein folding can also be assessed with the self-cleaving 2A peptide. This peptide, encoded by certain viral genes, is unique in its ability to self-cleave. When the ribosome translates the 2A peptide sequence from the mRNA, the resulting peptide interacts with the PET, triggering a conformational change that allows the release of the 2A peptide from the growing polypeptide chain and its independent exit from the ribosome. This short sequence (18-19 aa) is highly conserved among picornaviruses and is very sensitive to substitution (38).

We investigated whether the loss of Nm in the Δ4^PET^ strain affected the ability of 2A peptides to undergo efficient stop-carry on recoding. In the reporter system used (Fig 7A), 2A was sandwiched between a ubiquitin-arginine sequence (UBI-R) and the *ADE2* coding sequence (39). The removal of the UBI sequence leaves an arginine residue at the N-terminus of the 2A-Ade2 peptide, driving its degradation through the N-end rule pathway. Efficient 2A peptide hydrolysis allows the expression of a functional Ade2p protein devoid of the N-terminal arginine, thereby conferring adenine prototrophy and a red-to-white color change on the *ade2* mutant strain. The efficiency of 2A hydrolysis can be modulated through the use of 2A mutants. We hypothesized that changes in methylation status within the PET might have a positive or negative effect on self-cleavage of the 2A peptide. We, thus, tested mutants of the 2A peptide known to have partially affected activity (31 to 56% wild type levels) for the 15I, 14Q and 16H substitutions (40). The wild-type sequence and a sequence with a mutation of the terminal proline residue (19A, see Fig. 7*A*) essential for hydrolysis were used as the positive and negative controls, respectively. In both the control and mutant strains, the WT sequence of 2A conferred a light pink color on the cells, indicating that there was sufficient self-cleavage for the adenine biosynthetic pathway to be functional (Fig. 7*B*). By contrast, when the proline (WT) residue was replaced by an alanine (19A), no cleavage was observed and both strains were red due to the accumulation of a red intermediate of adenine biosynthesis. However, differences were observed for all three substitution mutants of the 2A peptide sequence, with a much more marked red color for the mutant ι14^PET^ strain than for the control. Thus, the self-cleaving activity of the 2A peptide is less efficient when the PET lacks the seven methylations investigated here, confirming the general defect of PET organization.

**Figure 7.**
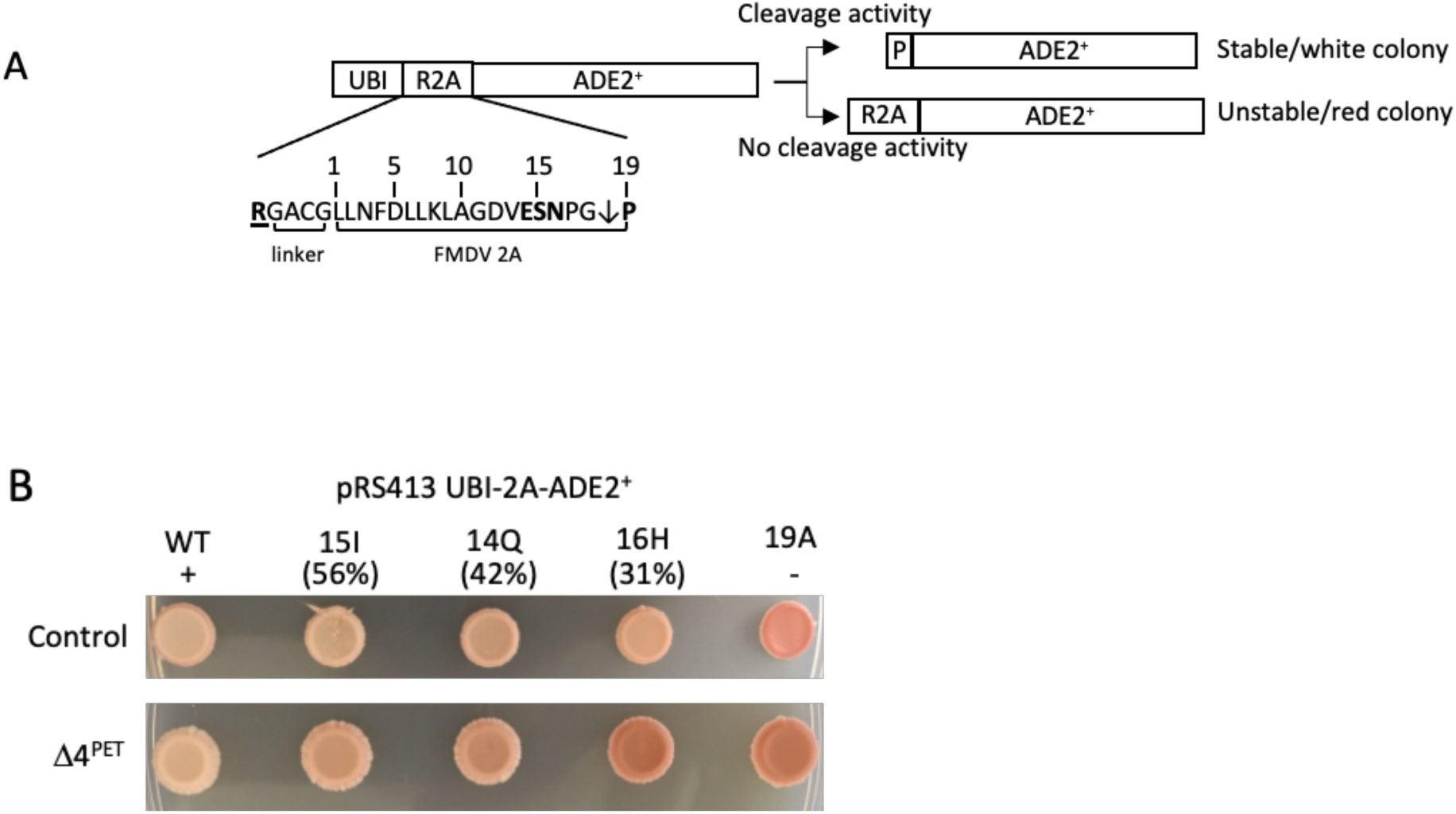
2A-based assay. (*A*) Structure of the WT reporter gene. The *ADE2^+^* coding sequence is fused to the UBI-Arg (R) sequence at its N-terminus, the two sequences being separated by the 2A self-cleaving peptide. The arginine residue responsible for protein instability is shown in bold and underlined. The amino acids of the 2A peptide are numbered and the cleavage site is indicated by an arrow between the terminal gly and pro residues. (*B*) Color assay. *Ade2* mutant strains with or without 4 snoRNA genes targeting modifications within the PET (1′4^PET^) were transformed with the reporter plasmid containing various [UBI-2A-ADE2] sequences. The WT is the positive control and 19A is the negative control. S15I, E14Q and N16H (in bold) present substitutions within the 2A sequence that alter its cleavage activity (%activity in the *in vitro* assay is shown in brackets). Strains were spotted onto a minimal SD medium supplemented with CSM and limited amounts of adenine and cultured for 3 days at 30°C.

## Discussion

Studies analysing the effects of rRNA modifications were done according to two different approaches. The first focused on the effects of an absence of the modifications clustering around functional sites of the ribosome, regardless of the real possibility of fluctuations of their status. The second was based on the partial depletion of either the essential enzymes dyskerin and fibrillarin, resulting in a lack of modifications at several sites within the ribosomes according to the capacity of these enzymes to generate variation but regardless of the site of the modifications. Here, we combined these two approaches to explore the potential of ribosomes to adapt translation through rRNA modification status. We chose *S. cerevisiae* as the model for this study due to its extraordinary amenability to genetic manipulations and the conservation of complex molecular processes between this yeast and humans (41). We found that 28% of the sites displaying a marked decrease in methylation upon fibrillarin depletion were conserved between this yeast and humans. This percentage is lower than the overall level of modification conservation of 50% between these two organisms (15, 17). Assuming that rRNA modifications fine-tune translation in response to environmental signals, then human cells — with their intricate cell-to-cell interactions and communication pathways crucial for tissue specificity and body development — would be expected to have a greater capacity to modulate modification status at a larger number of sites (42).

We then explored the impact of a complete loss of the variable modifications of the PET or the E site of the ribosome. We found that an absence of naturally occurring 2’-*O*-methylation at positions sensitive to fibrillarin knockdown had little or no effect on growth or on overall translation rates. This finding may reflect the fact that the manipulations were performed in precisely controlled laboratory conditions very different from those encountered by yeast cells in their natural environment, which is less demanding in terms of the need for adaptive (43). Marcel and coworkers explored the heterogeneity of methylation status in 195 primary human breast tumors and classified the 106 rRNA 2’-*O*-methylation sites into two groups: “stable” and “variable”. They then measured the proportion of sites in the “variable” category restricted to human ribosome and the proportion common to humans and yeast. They also found a smaller proportion of sites common to both yeast and humans than of sites specific to humans. They concluded that there was a link between modulation of the level of methylation of rRNA sites and the complexity of gene expression regulation (22).

Conversely, the introduction of several 2’-*O*-methylations at sites specific to human ribosomes on the intersubunit surface of the 60S had a major impact on yeast growth and translation generally, although the addition of a single 2’-*O*-methylation was without consequence. This approach has been used previously as an alternative to rRNA mutagenesis for targeting universally unmodified nucleotides within functionally important regions of the ribosome (44, 45). Methylation at positions G1927, G2240, C2304 and G2440 generally seems to be deleterious, suggesting a mutational effect. The interface between ribosomal subunits is highly conserved, with very few expansion segments and eukaryote-specific proteins (46). The association of the two subunits to form the 80S ribosome is crucial for translation. This association is promoted by multiple highly conserved contacts involving both rRNA and proteins and known as intersubunit bridges (47). The interface between the two subunits is also in contact with key components such as mRNA, tRNA and translational factors. The various interactions are dynamic and support the movements of intersubunits observed at all steps of translation, especially during translocation (review in (48)). A substantial number of modifications to eukaryotic rRNAs occur at the interface of the ribosomal subunits, where many rRNA segments participate in intersubunit bridge formation, suggesting that modifications to these bridge regions may influence the translation process. However, these findings raise questions about the reasons for such differences in 2’-*O*-methylation between the ribosome interfaces of yeast and humans and for the major effects of the introduction of these methylations on overall translation rates. One key finding of this study was the increase in CrPV initiation efficiency in the presence of at least 3 Nm (Gm1927+Gm2240+Cm2304 or Gm1927+Cm2304+Gm2440). The introduction of 2’-*O*-methylations at positions equivalent to those in the human ribosome, thus, confers enhanced CrPV IRES initiation activity on the yeast ribosome. This finding is consistent with the demonstration by Basu and coworkers that the activation of human c-srv IRES — a CrPV-like IRES — is specifically dependent on 60S methylation (49). Conversely, human *FBL* knockdown decreases CrPV IRES activity both *in vivo* and *in vitro* (25). However, neither of these effects could be attributed to specific positions. Structurally, helix 69 is known to interact with the D loop of P-site tRNA and has been reported to be in contact with the CrPV IRES (50). Am2256 (snR63) — one of the two variable 2’-*O*-methylations conserved between yeasts and humans, together with Cm2197 (snR76) — is part of helix 69, a component of intersubunit bridge B2a.

Interestingly, Metge and coworkers have shown that although the level of 2’-*O*-methylation at position A3739 (yeast A2256) is similar in normoxia and hypoxia, ribosomes bound to VEGF-IRES in hypoxia are specifically hypermethylated at this position. Thus, ribosomes with Am3739 are specifically associated with VEGF-IRES, confirming that specific methylation at the subunit interface of 60S promotes IRES binding (51). In addition, CrPV IRES internal loop L1.1 interacts with RPL1 and helices H76 and H77 of the 25S rRNA (17115051). The methylation of G2440, which lies within helix 76, could interfere with IRES activity. Yeast has no cellular IRES and viral IRES activity has, to date, been reported for only a few sequences (CrPV and HiPV). It could therefore be argued that if the 2’-*O*-methylation at these positions is involved in the efficiency of IRES initiation, then a modulation of modification status might be beneficial in human cells but dispensable in yeast cells.

The CrPV-IRES forms three pseudoknots (PKI-III) spanning all three tRNA binding sites of the ribosome, establishing many contacts with these three regions at the rRNA and ribosomal protein levels (50). Remarkably, the CrPV IRES undergoes a dynamic conformational change after two pseudotranslocation cycles, with the PKI domain in the ribosomal E site flipping from an anticodon-codon mimic (a conformation adopted when PKI is located within the A-site during initiation or within the P-site after the first pseudotranslocation step) to an acceptor stem of a canonical E-site tRNA mimic on the ribosome (52, 53). This change in conformation alters the interaction of the IRES with its environment, bringing the PKI closer to 25S rRNA helix 68 (H68). Given that snR47 drives 25S Am2220 2’-*O*-methylation at the apical tip of H68, we hypothesized that it might interfere with IRES activity. The results obtained with our dual reporter system showed there to be no direct effect of modifying this nucleotide.

Along similar lines, we found that 25S-U2421 was close to eL42-lys55 and eL42-Pro56 (rpl42) (Fig. 8), eL42-lys55 being the substrate of the ribosomal protein methyltransferase Rkm4 (54). If lys55 methylation is lost or Pro56 mutation occurs, yeast strains display cycloheximide-sensitive and cycloheximide-resistant phenotypes, respectively, attributable to the affinity of binding to the antibiotic (54–57). We assessed the cycloheximide tolerance of a strain lacking Snr78 and, consequently, Um2421. Remarkably, clear susceptibility potentially due to the same mechanism of action was observed in the absence of this Nm modification. Indeed, in bacteria, rRNA methylation is a natural and widespread mechanism preventing drug binding and favoring the acquisition of resistance to ribosome-targeting antibiotics (58).

**Figure 8.**
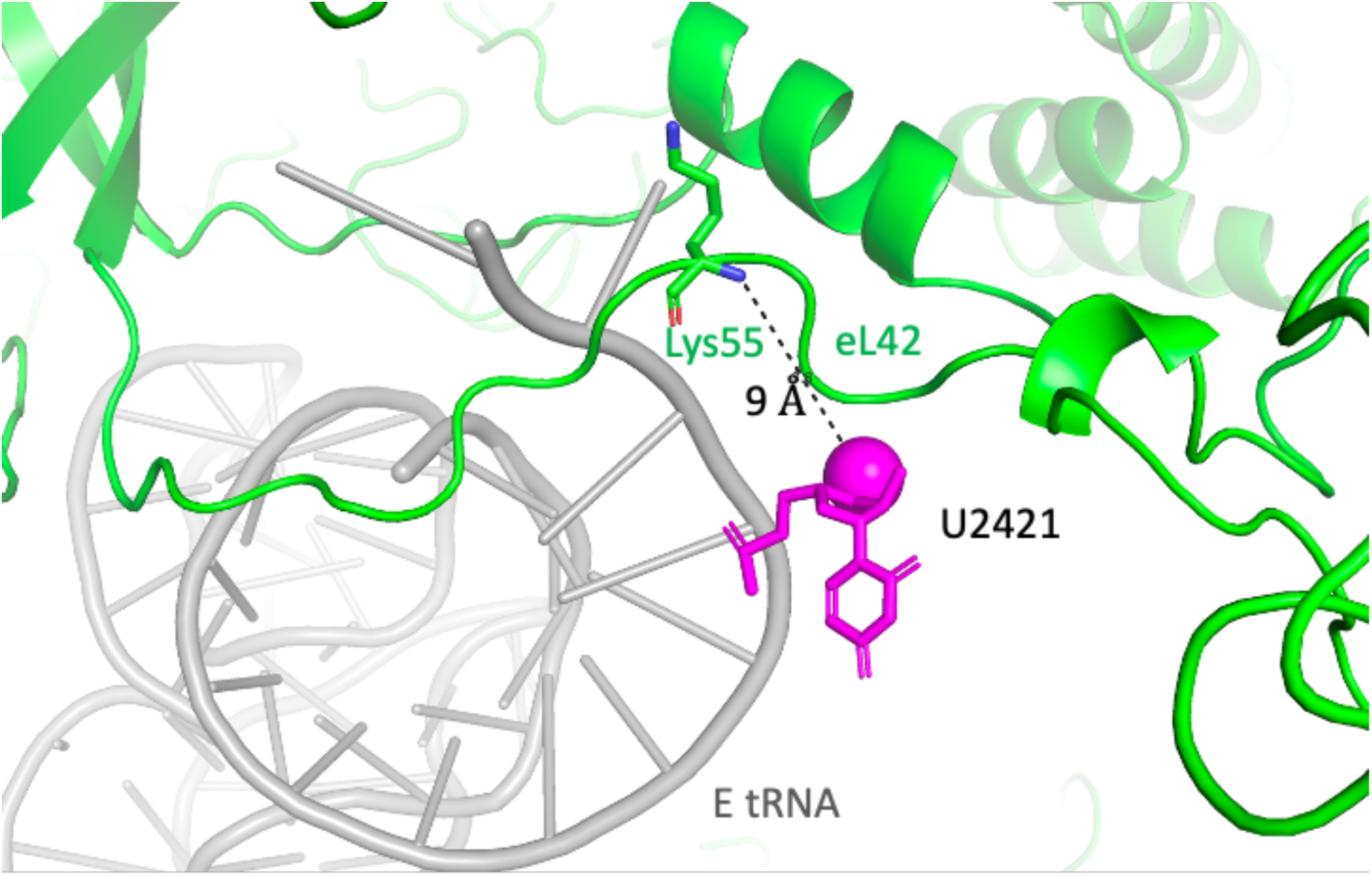
The Um2421 modification is located in close proximity to the Lys55 residue of eL42. Three-dimensional representation of the rRNA base of interest mapped onto the *S. cerevisiae* ribosome. Modified nucleotides are shown in magenta, with the methylated carbon depicted as a sphere, the eL42 peptide chain shown in green with Lys55 highlighted, and the E-site tRNA in gray.

The PET is a long channel that spans the 60S subunit from the PTC to the outer surface and through which the nascent polypeptide passes. It involves five of the six domains of the 25S rRNA and the RPs uL4 (rpL4), uL22 (rpL17), and eL39 (rpL39) (46). One of these proteins, uL4, plays a key role in shaping the two constriction sites required for correct translation. It has become clear that the PET is not just a simple exit route for the nascent peptide. It is also involved in translation control and peptide structure through its interactions with rRNA residues and ribosomal proteins (59, 60). In parallel, PET, as one of the functionally important regions of the ribosome, has been shown to be rich in nucleotide modifications (25). This raises the possibility that 2’-*O*-methylations modulate PET activity. We tested this hypothesis by measuring the self-cleaving activity of the 2A peptide, which was decreased in the absence of modifications susceptible to FBL knockdown. Careful examination of the location of these modifications in this region showed that three of these modifications were located close to the universal ribosomal protein uL4 and the β-hairpin protruding into the tunnel (Fig. 9). Gm805 and Cm1437 are directly opposite and close to uL4 Arg7 and Arg73, respectively, near the top of the tentacle domain and on the tunnel surface, whereas Cm663 is close to uL4 Arg107 at the bottom of the β-hairpin. Am649 and Cm650 are located opposite Gm805 and Cm1437 near the surface of the exit tunnel. Changes in methylation within the rRNA in close contact with the tentacle domain of uL4 in our mutant may, therefore, affect the conformation of the exit tunnel, preventing the formation of the α-helix and the reverse turn of the 2A peptide required for correct hydrolysis of the peptidyl–tRNA^Gly^ ester bond and release of the Ade2 protein.

**Figure 9.**
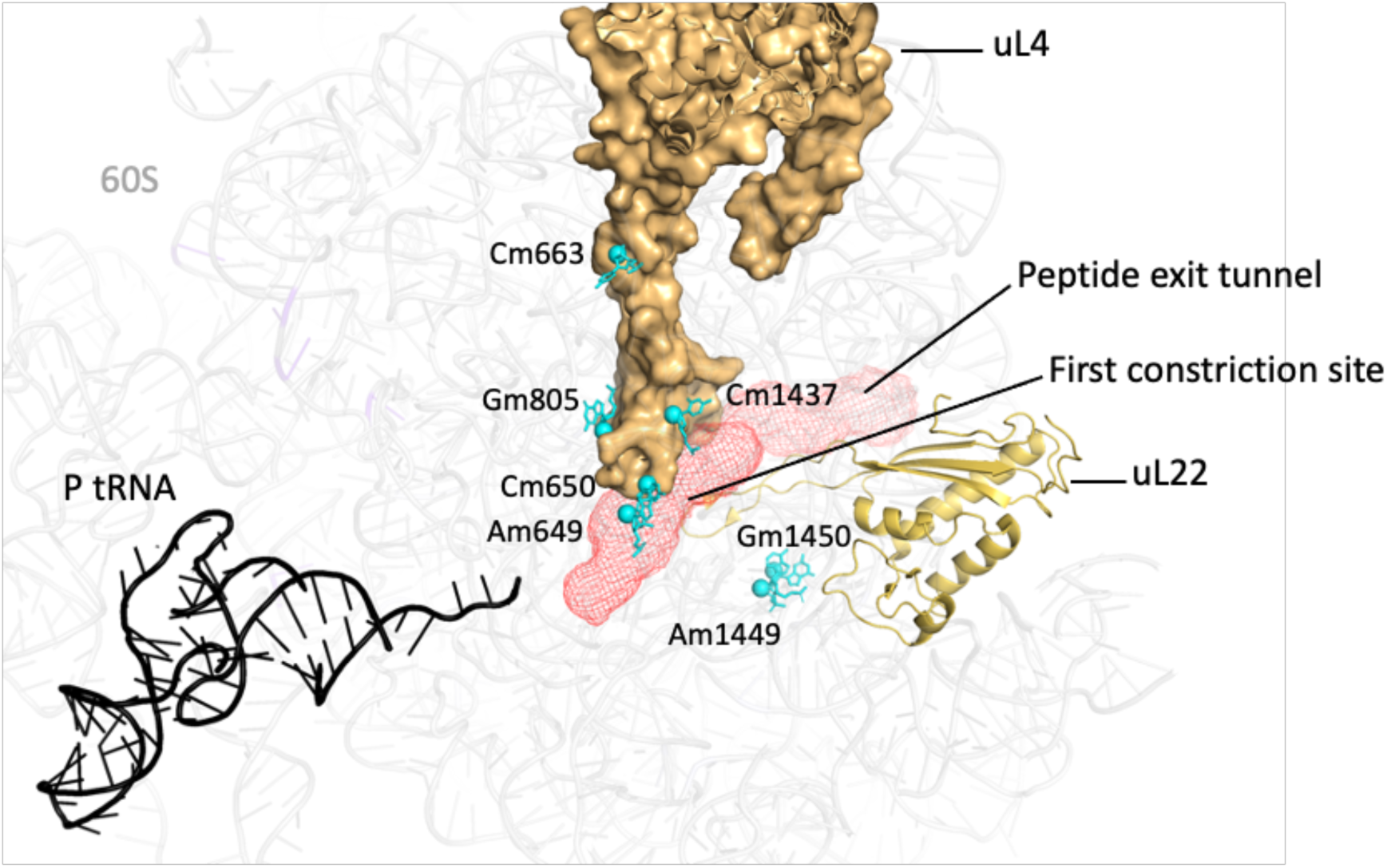
Tertiary structure of the ribosome around the peptide exit tunnel. CryoEM structure of the yeast 60S subunit (PDB 3J78). The peptide exit tunnel coordinates (in red) were extracted as described by K. Dao Duc et al. (30805621, 36305870). The two proteins involved in the first constriction site (uL4 and uL22) are depicted in yellow; The P-site tRNA is shown in black and the nucleotides lacking methylation (sphere) in the 1′4^PET^ strain shown in turquoise.

This study reveals a broader cellular role for rRNA 2’-*O* methylation beyond simply stabilizing or folding rRNA. By demonstrating the influence of 2’-*O* methylation on both IRES-dependent translation initiation and the function of the peptide exit tunnel, this study highlights its multifaceted role in regulating protein synthesis and protein folding. These findings provide support for the notion that chemical modifications of the ribosome can regulate gene expression at the translational level.

## EXPERIMENTAL PROCEDURES

### Yeast manipulation and standard molecular biology

Yeast strains lacking single or multiple snoRNA genes were constructed by sequential gene disruption (61) in strain FY1679-28C or FY1679-18B (*MAT*a and *MAT*α, respectively, *ura3-52 trp1β63 leu2β1 his3β200 GAL2*) (62). Strain Δ4^PET^ contains plasmids pRS416-*ASC1*orf (29) and pCM189-*EFB1*orf (63) with or without pRS413-ACT/snR24 (29). The target nucleotides were methylated as previous described (45). Briefly, a new plasmid-encoded guide snoRNA was derived from the natural snR38 gene. This novel methylation guide was expressed under the control of the *GAL1* promoter. The artificial guide sequence was inserted by a PCR-based strategy into plasmid pBL150 using custom oligonucleotide tacaaatatcaacatatgcattcagNNNNNNNNNNNNNtttgataaaattttttttcatc with CTTCGGGATAAGG resulting in pBL(G1927), GCGGGTAAACGGC resulting in pBL(G2240), GTGACGCGCATGA resulting in pBL(C2304) and TGAAGAGACATAG resulting in pBL(G2440), with the targets indicated in brackets. The parental snR38, which targets G2811 in 25S rRNA, was used as control (pBL152). Methylation of the four targeted nucleotides was confirmed by RiboMethSeq analysis (15). Cells were grown in complete YPD medium (1% yeast extract, 2% peptone, 2% dextrose) or in minimal synthetic defined (SD) medium containing glucose (2% glucose, 0.7% yeast nitrogen base) or galactose (2% galactose, 0.7% yeast nitrogen base) and supplemented with Complete Supplement Mixture lacking the appropriate amino acids or bases (MP Biomedicals) according to standard procedures. The strains used in this study are listed in Table 2.

### Computational analysis of ribosome structure and the peptide exit tunnel

The crystal structure of the *S. cerevisiae* ribosome was visualized and analyzed with PyMOL DeLano (http://pymol.org/) and PDB 3J78. Peptide exit tunnel coordinates were obtained from Khanh Dao Duc (64, 65).

### Ribosome–polysome profile analysis

Cell extracts were prepared and ribosome–polysome profiles were analyzed as previously described (66). For polysome/monosome (P/M) ratio quantification, the areas under the 80S and polysome peaks were quantified with ImageJ. The mean and SD for at least three biological replicates were calculated.

### Antibiotic susceptibility test

Wild-type and mutant cells were grown in YPD or YNB to an OD_600_ of 1.0. Tenfold dilutions from 1 to 10^-5^ were prepared. We spotted10 µl of dilutions from 10^-3^ to 10^-5^ on YPD or YNB plates containing the antibodies indicated. The following concentrations were used: 0.5 μg/ml for cycloheximide and 50, 100, 250 and 500 ng/ml) for canavanine. Arginine was omitted from the medium, when necessary, to ensure the efficient import of canavanine into cells. Plates were incubated at 30°C for 3 days.

### Growth studies

Growth was measured in triplicate in minimal medium supplemented with Complete Supplement Mixture (CSM) lacking the appropriate amino acids or bases (MP Biomedicals). Growth was monitored for 24 to 48 hours at 30°C by measuring the OD at 600 nm with a Tecan Infinite plate reader.

### Quantification of translation accuracy

Strains were transformed with a set of dual reporter plasmids carrying an upstream *lacZ* and a downstream firefly luciferase reading frame separated by an in-frame stop codon in the tobacco mosaic virus (TMV) readthrough context (CCA GCA GGA ACA CAA <u>TAG</u> CAA TTA CAG TGG), the -1 frameshift sequence of the infectious bronchitis virus (IBV) (TATTTAAACGGGTACGGGGTATCAGTCAGCTCGGCTGGTACCCCTTGCAAAGCGAGCCTCA), the -1 frameshift sequence of the bovine leukemia virus (BLV) (AAATCAAAAAACTAATAGAGGGGGGACTTAGCGCCCCCCAAACCGTA), the +1 frameshift of the *S. cerevisiae EST3* gene (CAAATACTTAGTTGAGTTTTCC) or the control sequence (CCAGCAGGAACACAACAGCAATTACAGTGG). Beta-galactosidase activity and luciferase activity were estimated as previously described (67). The firefly luciferase activity measured reflects translation error, whereas the β-galactosidase activity serves as an internal normalization control. Error efficiency was calculated as the luciferase/β-galactosidase activity ratio expressed as a percentage of that for an in-frame fusion construct. The “fold WT” value corresponds to the mutant versus wild-type efficiency ratio. The data shown are the mean ± SEM (*n* = 5). Statistical significance was determined by calculating *p*-values in *t*-tests.

### Quantification of IRES initiation

For the quantification of IRES initiation in yeast, we used the pRST215 dicistronic reporter plasmid, in which the CrPV IGR IRES controls the expression of firefly luciferase whereas that of the *Renilla* luciferase is controlled by the cap-dependent pathway (68). A second type IV IRES, HiPV IRES, was inserted into the dicistronic vector pTH779 (69) and tested in cases of negative results with pRST215.

Yeast transformed with *URA3* dicistronic luciferase plasmids were prepared for luciferase assays by overnight culture in selective media. We harvest 1 OD_600_ unit of yeast in mid-exponential growth phase and resuspended it in 100 μl Passive Lysis Buffer (Promega) with mixing for 2 minutes. Luciferase assays were performed with the Dual-Luciferase Reporter Assay System (Promega) according to the manufacturer’s protocols. Activity was measured with a TECAN luminometer. The data shown are the mean ± SEM (*n* = 5). Statistical significance was determined by calculating the *p*-value in a *t*-test.

### Translation assay

We quantified the amounts of newly synthesized protein within yeast cells by measuring the incorporation of L-HPG, an amino acid analog. L-HPG was detected via an Alexa Fluor 488 moiety attached by click chemistry (Click-iT® HPG Alexa Fluor® Protein Synthesis Assay Kits Catalog nos. C10428, C10429, Invitrogen). The level of fluorescence measured by cytometry reflects the rate of translation. This experiment was performed with a protocol described in a previous study (70).

### Analysis of rRNA methylation by RiboMethSeq

Total RNA (100 ng) from yeast *S. cerevisiae* cells was subjected to alkaline hydrolysis in 50 mM sodium bicarbonate buffer pH 9.2 for 14 min at 96°C. The reaction was stopped by precipitation in ethanol in the presence of 3M sodium acetate, pH 5.2 and GlycoBlue. The precipitate was collected by centrifugation and the pellet was washed with 80% ethanol and resuspended in nuclease-free water. RNA fragments were end-repaired as previously described (15) and purified with the RNeasy MinElute Cleanup kit according to the manufacturer’s recommendations except that 675 µl of 96% ethanol was used for RNA binding. Elution was performed in 19 µL nuclease-free water. RNA fragments were used to generate a library with the NEBNext R© Small RNA Library kit (NEB ref E7330S, UK), according to the manufacturer’s recommendations. The resulting DNA library was quantified with a fluorometer (Qubit 2.0 fluorometer, Invitrogen, USA) and subjected to a qualitative analysis on a high-sensitivity DNA chip on an Agilent Bioanalyzer 2100. Libraries were multiplexed and subjected to high-throughput sequencing on an Illumina HiSeq1000 instrument in 50 bp single-end read mode.

Sequencing reads were trimmed with Trimmomatic v0.32, and the trimmed reads were aligned with the *S. cerevisiae* rRNA reference sequence and its protection profile was calculated as previously described (15). RiboMethSeq scores were calculated for all rRNA positions to ensure the absence of unexpected Nm residues.

### Availability of datasets

The raw RiboMethSeq data are available from the European Nucleotide Archive under accession number PRJEB77114.

## ACKNOWLEDGMENTS

We thank J.D. Brown for kindly providing the Ubi-R2A-ADE2 vectors, and V. Marcel and F. Catez for providing IRES-containing vectors and for discussions. We are very grateful to K. Dao Duc for calculating the tunnel structure from the 3J78 pdb file. This project was funded by the *Agence Nationale pour la Recherche* (ACTIMETH 19-CE12-0004-02).

